# Revealing mechanisms of infectious disease transmission through empirical contact networks

**DOI:** 10.1101/169573

**Authors:** Pratha Sah, Michael Otterstatter, Stephan T. Leu, Sivan Leviyang, Shweta Bansal

## Abstract

The spread of pathogens fundamentally depends on the underlying contacts between individuals. Modeling infectious disease dynamics through contact networks is sometimes challenging, however, due to a limited understanding of pathogen transmission routes and infectivity. We developed a novel tool, INoDS (Identifying Network models of infectious Disease Spread) that estimates the predictive power of empirical contact networks to explain observed patterns of infectious disease spread. We show that our method is robust to partially sampled contact networks, incomplete disease information, and enables hypothesis testing on transmission mechanisms. We demonstrate the applicability of our method in two host-pathogen systems: *Crithidia bombi* in bumble bee colonies and Salmonella in wild Australian sleepy lizard populations. The performance of INoDS in synthetic and complex empirical systems highlights its role in identifying transmission pathways of novel or neglected pathogens, as an alternative approach to laboratory transmission experiments, and overcoming common data-collection constraints.

## Introduction

Host contacts, whether direct or indirect, play a fundamental role in the spread of infectious diseases (*Newman, 2002; Rohani et al., 2010; Bansal et al., 2007; Sah et al., 2017a*). Traditional epidemiological models make assumptions of homogeneous social structure and mixing among hosts which can yield unreliable predictions of infectious disease spread (*Shirley and Rushton, 2005; Volz and Meyers, 2007; Bansal et al., 2007; Chen et al., 2014*). Network approaches to modeling the spread of infectious diseases provide an alternative by explicitly incorporating host interactions that mediate pathogen transmission. Formally, in a contact network model, individuals are represented as nodes, and an edge between two nodes represents an interaction that has the potential to transmit infection. Constructing a complete contact network model requires (*i*) knowledge about the transmission routes of a pathogen, (*ii*) a sampling of all individuals in a population, and (*iii*) a sampling of all disease-causing interactions among the sampled individuals. In addition, accuracy of disease predictions depends on the quantification of the epidemiological characteristics of the pathogen, including the rate of pathogen transmission given a disease-causing contact between two individuals, and the rate of recovery of infected individuals.

The use of modern technology in recent years, including RFID, GPS, radio tags, proximity loggers and automated video tracking has enabled the collection of detailed movement and contact data, making network modeling feasible. Despite the technology, logistical and financial constraints still prevent data collection on all individuals and their social contacts (*Welch et al., 2011; Cross et al., 2012; Godfrey, 2013; Krause et al., 2013; Farine and Whitehead, 2015; Silk et al., 2015*). More importantly, limited knowledge about a host-pathogen system makes it challenging to identify the mode of infection transmission, define the relevant disease-causing contacts between individuals, and measure per-contact rate of infection transmission (*Craft and Caillaud, 2011; White et al., 2015; Manlove et al., 2017*). Laboratory techniques of unraveling transmission mechanisms usually take years to resolve (*Velthuis et al., 2007; Aiello et al., 2016; Antonovics et al., 2017*). Defining accurate contact networks underlying infection transmission in human infectious disease has been far from trivial (*Bansal et al., 2007; Pellis et al., 2014; Eames et al., 2015*). For animal infectious disease, limited information on host behavior and the epidemiological characteristics of the spreading pathogen makes it particularly difficult to define a precise contact network, which has severely limited the scope of network modeling in animal and wildlife epidemiology (*Craft and Caillaud, 2011; Craft, 2015*).

Lack of knowledge about disease transmission mechanisms has prompted the use of several indirect approaches to identify the link between social structure and disease spread. A popular approach has been to explore the association between social network position (usually quantified as network degree) of an individual and its risk of acquiring infection (*Godfrey et al., 2009, 2010; Leu et al., 2010; MacIntosh et al., 2012*). Another approach is to use proxy behaviors, such as movement, spatial proximity or home-range overlap, to measure direct and indirect contact networks occurring between individuals (*Danon et al., 2011; Hamede et al., 2009; Fenner et al., 2011).* A recent approach, called the *k*-test procedure, explores a direct association between infectious disease spread and a contact network by comparing the number of infectious contacts of infected cases to that of uninfected cases (*VanderWaal et al., 2016*). However, several challenges remain in identifying the underlying contact networks of infection spread that are not addressed by these approaches. First, it is often unclear how contact intensity (e.g. duration, frequency, distance) relate to the risk of infection transfer unless validated by transmission experiments (*Aiello et al., 2016)*. Furthermore, the role of weak ties (i.e., low intensity contacts) in pathogen transfer is ambiguous (*Pellis et al., 2014; Sah et al., 2017b*). The interaction network of any social group will appear as a fully connected network if monitored for a long period of time. As fully-connected contact networks rarely reflect the dynamics of infectious disease spread through a host population, one may ask whether weak ties can be ignored, or what constitutes an appropriate intensity threshold below which interactions are epidemiologically irrelevant? Second, many previous approaches ignore the dynamic nature of host contacts. The formation and dissolution of contacts over time is crucial in determining the order in which contacts occur, which in turn regulates the spread of infectious diseases through host networks (*Bansal et al., 2010; Fefferman and Ng, 2007; Farine, 2017*). Finally, none of the existing approaches allow direct comparison of competing hypotheses about disease transmission mechanisms which may generate distinct contact patterns and consequently different contact network models.

All of these challenges demand an approach that can allow direct comparison between competing contact network models while taking into account the dynamics of host interactions and data-constraints of network sampling. In this study, we introduce a computational tool called INoDS (*I*dentifying *N*etwork models of infectious .Disease *S*pread) that quantifies the predictive power of empirical contact networks in explaining infectious disease spread, and enables comparison of competing hypotheses about transmission mechanisms of infectious diseases. Our tool can also infer the per-contact transmission rate of various infectious disease types (SI, SIS, and SIR), and can be easily extended to incorporate other complex models of disease spread. The INoDS tool provides inference on dynamic and static contact networks, and is robust to common forms of missing data. Using two empirical datasets, we highlight the two-fold application of our approach - (*i*) to identify whether observed patterns of infectious disease spread are likely given an empirical contact network, and (*ii*) to identify transmission routes, the role of the contact intensity, and the per contact transmission rate of a host-pathogen system. The epidemiological mechanisms of infection transmission identified by INoDS can therefore provide invaluable insights during implementation of immediate disease control measures in the event of an epidemic outbreak.

## Results

The primary purpose of INoDS is to assess whether an empirical contact network is likely to generate an observed infection time-series from a particular host population. INoDS also provides epidemiological insights into the spreading pathogen by estimating the per-contact rate of transmission. In practice, the structure of a contact network model depends on the mode of infection transmission, and is sensitive to the amount of missing data on nodes and edges. The tool therefore treats empirically collected contact network models as network hypotheses, and facilitates hypothesis testing between different contact networks. The INoDS algorithm follows a three step procedure (Figure 1). First, the tool estimates a per-contact pathogen transmission rate (*β*) and an error parameter (*ε*). The *β* parameter quantifies the rate of pathogen transmission through each edge of the contact network, and the *ε* parameter quantifies components of infection transmission that are unexplained by the edge connections of the contact network. In the second step, the likelihood of the observed infection time-series under the network hypothesis and pathogen transmission rate is compared to the null likelihood distribution based on an ensemble of randomized networks. The randomized networks are generated by permuting the edge connections of the network hypothesis, while controlling the number of nodes and edges present. We consider the network hypothesis to have high predictive power if the likelihood of thr infection time-series given the hypothesis is higher than the null likelihoods at the 5% significance level. In the final step, the marginal (Bayesian) evidence is calculated for the network hypothesis, which can be used to perform model selection between multiple network hypotheses.

**Figure 1.**
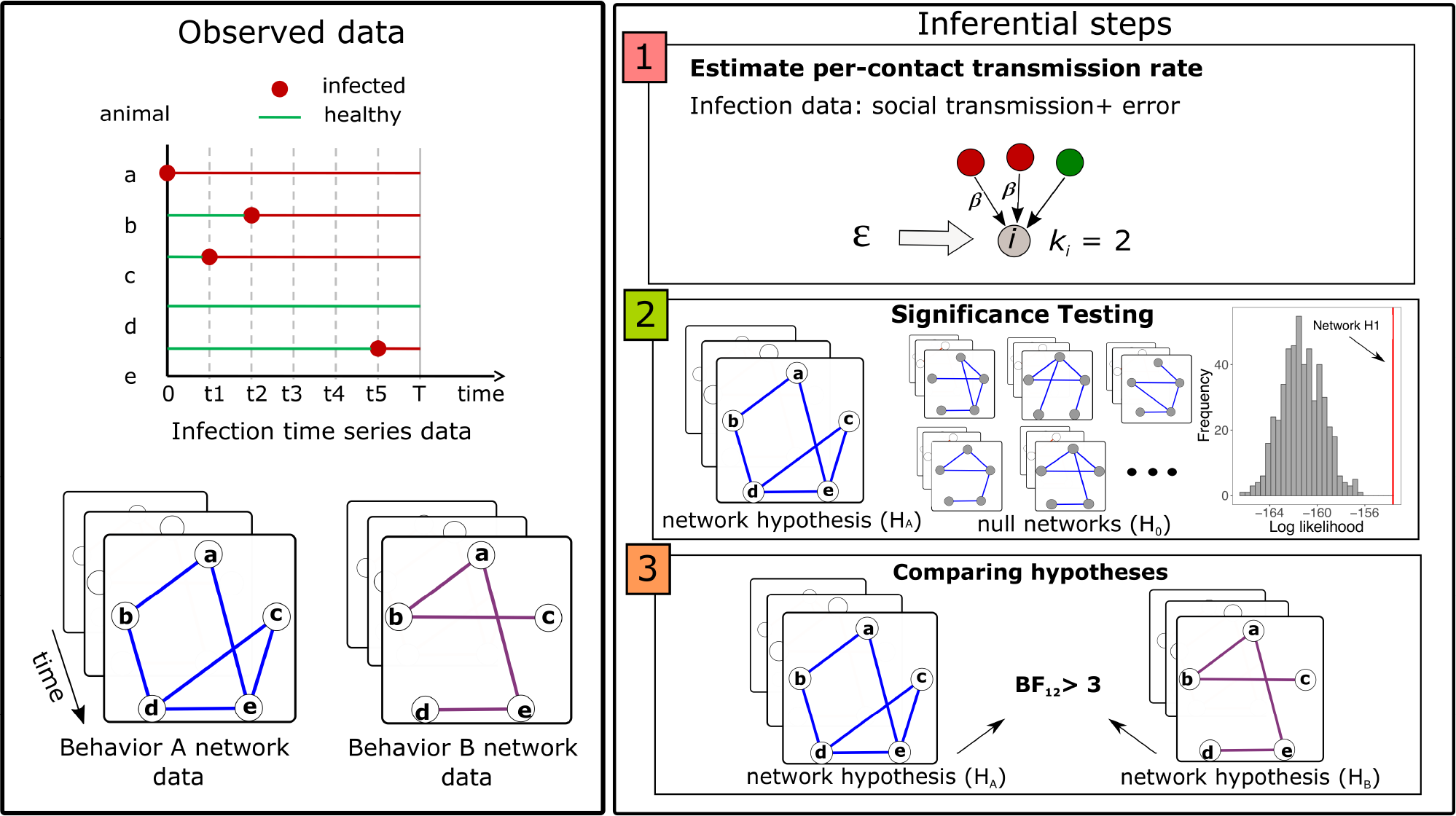
A schematic of our algorithm. **Observed data**: INoDS utilizes an observed infection time-series to estimate evidence towards a static or dynamic contact network hypothesis (or hypotheses) using a three step procedure. Shown here is an example of two competing contact network hypothesis based on different de1nitions of disease-causing contact (quanti1ed by behavior A and behavior B). **Inferential steps**: In the 1rst step, the tool estimates per-contact transmission rate parameter *β*, and an error parameter ε which captures the components of infection propagation unexplained by the edge connections of the network hypothesis. Second, the likelihood that the infectious disease spreads through the edge connections of the contact network hypothesis is compared to a distribution of likelihoods obtained from an ensemble of randomized networks. The predictive power of the empirical network hypothesis is considered to be high when its likelihood is higher than the null likelihood distribution at 5% signi1cance level. Third, the marginal likelihood for the contact network hypothesis is calculated, which is then used to perform model comparison (using Bayes Factor, BF) between multiple contact network hypotheses, wherever available.

In the sections that follow, we evaluate the accuracy of the tool in recovering the transmission parameter, *β*, and its robustness to missing data (missing individuals, missing contacts and missing infection cases). We further demonstrate the application of INoDS by using two empirical datasets: (*i*) spread of an intestinal pathogen in bumble bee colonies, and (*ii*) salmonella spread in Australian sleepy lizards.

## INoDS performance

We evaluated the performance of INoDS on multiple infection time-series data generated by performing numerical simulations of infection spread on a synthetic dynamic network, for a wide range of pathogen transmission rates.

We found that INoDS accurately estimates the true value of pathogen transmission rate, *β*, and the accuracy is independent of the spreading rate of the pathogen (Figure 2). The error parameter, *ε*, specified in the algorithm improves the estimate of transmission rate when either the network data or disease surveillance is incomplete (Appendix Figure 1). The estimated rate of pathogen transmission is therefore accurate even when substantial network or infection time-series data is missing (Appendix Figure 2). The expected value of *ε* is zero when all infection transmission events are explained by the edge connections in the contact network hypothesis (Figure 2). Values greater than zero, on the other hand, indicate unexplained transmission events due to either missing or inaccurate data (Appendix Figure 2).

**Figure 2.**
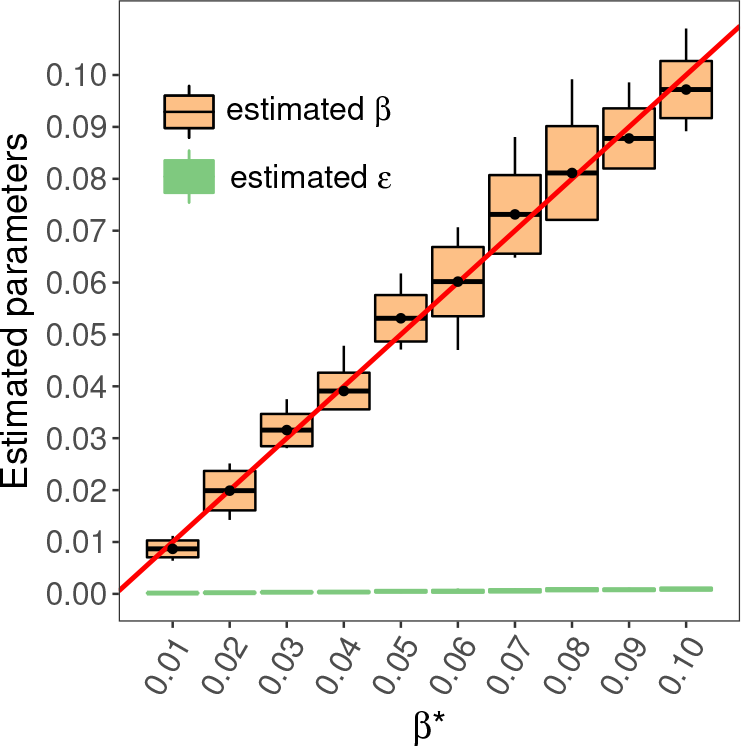
INoDS performance in recovering the per-contact pathogen transmission rate (*β*) for simulated infection time series under an susceptible-infected (SI) model. Each boxplot summarizes the results of 10 independent disease simulations; the black horizontal lines are the means of the estimated parameter values, the top and bottom black horizontal lines represent the standard deviation, and the tip of the black vertical line represents the maximum/minimum value. The solid red line represents one-to-one correspondence between the true value of the pathogen transmission rate (used to generate the simulated data), *β^*^*, and the *β* value estimated by INoDS. Since the simulations were performed on a known synthetic network, the expected value of the error parameter, *ε*, (represented by the green lines) is zero.

Next, we tested the performance of INoDS in establishing the epidemiological relevance of a hypothesized contact network against three potential sources of error in data-collection: (a) incomplete sampling of individuals in a population (missing nodes); (b) incomplete sampling of interactions between individuals (missing edges); and (c) infrequent health diagnosis of individuals (missing cases). The performance of the tool was quantified in terms of a true positive rate (i.e., the proportion of times an epidemiologically relevant contact network with missing data was correctly distinguished as statistically significant from an ensemble of randomized networks with the same amount of missing data) and a true negative rate (i.e., the proportion of times a network with the same degree distribution as the epidemiologically relevant contact network, but with randomized edge connections, was correctly classified as statistically insignificant). We found INoDS to be both sensitive and specific (with a high true positive and true negative rate) across a range of missing data scenarios (Figure 3). The true positive rate of the tool remains close to one even when as low as 10% of infection cases are documented (missing cases, Figure 3). For incompletely sampled contact networks, the true positive rate remains close to one when at least 50% nodes or 30% edges are documented.

**Figure 3.**
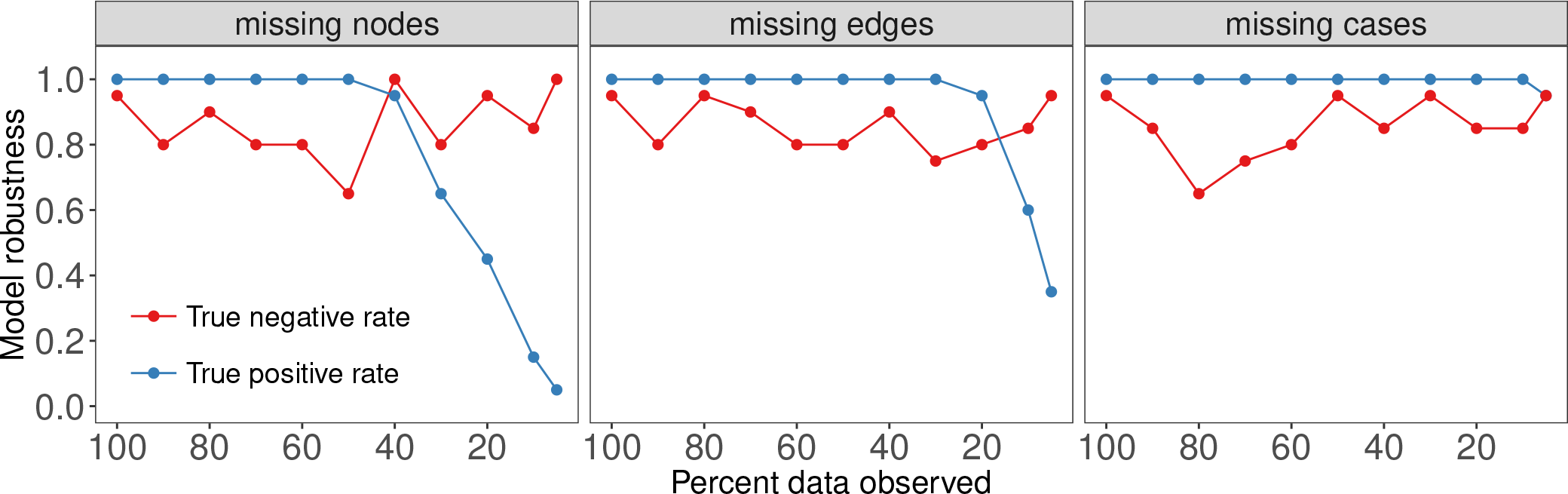
Robustness of INoDS in establishing the epidemiological significance of a hypothesized contact network under three common forms of missing data - missing nodes, missing edges and missing infection cases. True positive rate is calculated as the proportion of times (*n* = 20) the epidemiologically relevant (true) dynamic network was detected as statistically significant relative to a null distribution of randomized networks. True negative rate is calculated as the proportion of times (*n* = 20) a network with similar degree distribution as the epidemiologically relevant network, but randomized edge connections, was identified as statistically indistinguishable from the null distribution. The null distribution for a network hypothesis was generated by permuting all its edge connections, but preserving the number of nodes and edges.

The performance of INoDS in discriminating the epidemiological contact network from null network hypotheses also surpasses two previous approaches - the *k*-test procedure and the network position test (Figure 3 - figure supplement). In comparison to INoDS, the *k*-test and network position test are sensitive to all three types of missing data. The true positive rate of the fc-test declines with an increasing number of missing nodes, edges, or infection cases. Of the three approaches, the network position test has the lowest sensitivity (true positive rate). Since the fc-test procedure and network position test have been primarily used in the context of non-dynamic networks, we repeated this analysis with simulated disease data from a static synthetic network. Appendix Figure 4 demonstrates that even for observed networks that are not dynamic, INoDS has greater sensitivity and specificity than the *k*-test procedure and the network position test.

## Applications to empirical data-sets

We next demonstrate the application of INoDS to perform hypothesis testing on contact networks, identify transmission mechanisms and infer transmission rate using two empirical datasets. The first dataset is derived from the study by *Otterstatter and Thomson* (*2007*) that examined the spread of an intestinal pathogen (*Crithidia bombi*) within colonies of the social bumble bee, *Bombus impatiens*. The second dataset documents the spread of *Salmonella enterica* within two wild populations of Australian sleepy lizards *Tiliqua rugosa* *Bull et al.* (*2012*).

### Determining transmission mechanism and the role of contact intensity: case study of *Crithidia bombi* in bumble bees

*Otterstatter and Thomson (2007)* showed that the transmission of *C. bombi* infection in bumble bee colonies was associated with the frequency of contacts with infected nest-mates rather than the duration of contacts. However, the dynamic contact network models had a small number of nodes, and were fully connected (i.e., all individuals were connected to each other in the network) at all time steps. Because predictions of infection transmission is sensitive to the size and edge density of the contact network model, we extended the previous analysis by answering three specific questions: (1) Do physical contact networks have higher predictive power to explain the spread of *C. bombi* than random networks?, (2) Do the value of contact intensities (edge weights) matter in transmission?, and (3) Do weak ties between individuals contribute to infection transfer? To validate our tool, we performed analyses on two types of contact network models - those described by frequency of contacts and those described by duration of contacts - and compared the results with the findings reported in (*Otterstatter and Thomson, 2007*).

To answer the three questions, we constructed dynamic contact networks where edges represent close proximity between individuals. Since fully connected networks rarely describe the dynamics of infection spread, we sequentially removed edges with weights less than 10-50% of the highest edge weight to generate contact network hypotheses at different edge weight thresholds. Corresponding to the two types (frequency and duration) of weighted networks, unweighted contact networks were also constructed by replacing weighted edges in the thresholded weighted networks with binary edges (i.e., edges with an edge weight of one).

Figure 4 shows the estimates of pathogen transmission rate *β*, and error *ε* for the four types of contact network hypotheses at different edge weight thresholds. Only a subset of contact network hypotheses had statistically significant estimates of *β* (non faded bars). Two network hypotheses summarizing frequency of contacts - binary frequency network at 15% edge weight threshold and weighted frequency network at 5% edge weight threshold - demonstrated higher predictive power than an ensemble of null networks. Of the two, the weighted frequency network had slightly higher Bayesian evidence, although the binary frequency network was equally supported. In a separate colony (UN1), only the weighted frequency network at 5% edge weight threshold had higher predicted power compared to an ensemble of null networks (Appendix Figure 5). Together, our results therefore show that (1) contact networks capturing frequency (but not duration) of contacts have statistically high predictive power to explain the spread of *C. bombi* in bumble bee colonies, (2) the contact networks should be weighted, and (3) weak ties (i.e., edges with weights less than 5% of the highest weighted edge) are epidemiologically unimportant.

**Figure 4.**
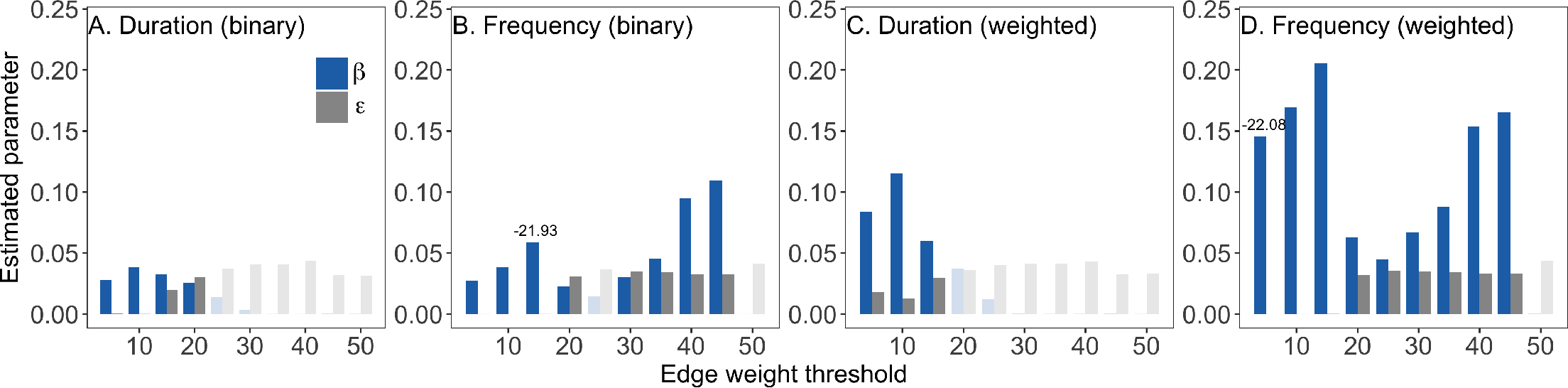
Identifying the contact network model of *Crithidia* spread in bumble bee colony (colony UN2). Edges in the contact network models represent physical interaction between the bees. Since the networks were fully connected, a series of 1ltered contact networks were constructed by removing weak weighted edges in the network. The x-axis represents the edge weight threshold that was used to remove weak edges in the network. Two types of edge weights were tested - duration and frequency of contacts. In addition, across all ranges of edge weight threshold, the weighted networks were converted to binary networks. The results shown are estimated values of the per contact rate transmission rate *β*, and estimated values of error *ε*, for the (A-B) the two types of binary network, (C) contact duration weighted network, (D) contact frequency weighted network. The faded bars correspond to networks where *β* parameter is statistically insigni1cant. Numbers above bars indicate the log Bayesian (marginal) evidence of the networks that were detected to have statistically signi1cant higher predictive power as compared to an ensemble of null networks (P < 0.05, corrected for multiple comparisons.

### Identifying transmission mechanisms with imperfect disease data: case study of *Salmonella enterica* Australian sleepy lizards

Spatial proximity is known to be an important factor in the transmission of *Salmonella enterica* within Australian sleepy lizard populations (*Bull et al., 2012*). However, it is not known whether the transmission risk increases with frequency of proximate encounters between infectious and susceptible lizards. We therefore tested two contact network hypotheses to explain the spread of salmonella at two sites of wild sleepy lizards populations. The first contact network hypothesis placed binary edges between lizards if they were ever within 14m distance from each other during a day (24 hours). We constructed the second contact network by assigning edge weights proportional to the number of times two lizards were recorded within 14m distance of each other during a day.

Because disease sampling was performed at regular fortnightly intervals, the true infection time (day) of individuals at both study sites was unknown. We therefore used a data augmentation method in INoDS (see Methods) to sample unobserved infection timings along with the per contact transmission rate, *β*, and error, *ε*. We found that the likelihood of salmonella infection spreading through the weighted contact network was significantly greater than the null expectation at both sites (Figure 5). Compared to unweighted networks, networks with edges weighted by contact frequency had higher marginal (Bayesian) evidence at both sites. This indicates that the occurrence of repeated contacts between two spatially proximate individuals, rather than just the presence of contact between individuals is important for Salmonella transmission.

**Figure 5.**
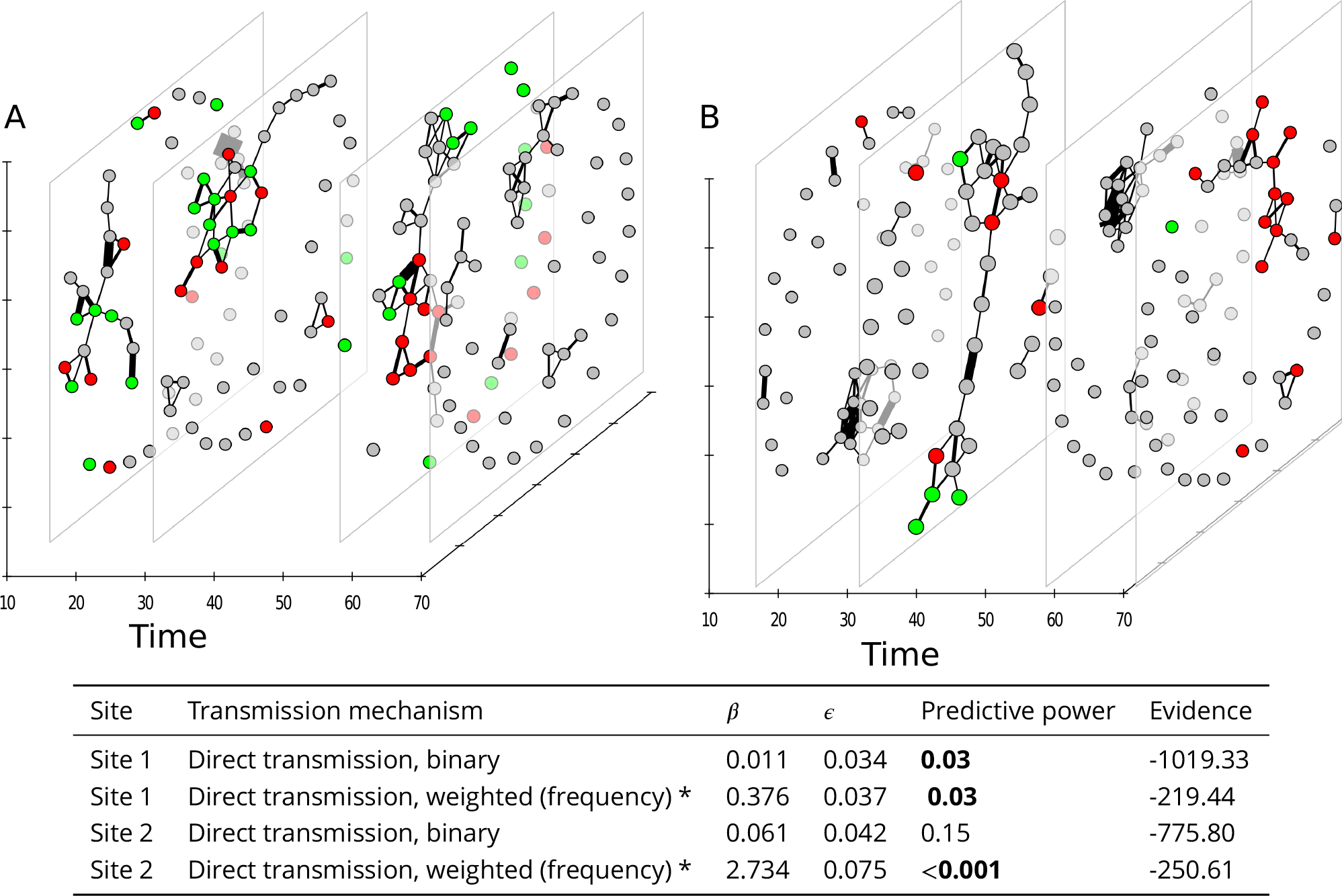
Identifying transmission mechanisms of Salmonella spread in Australian sleepy lizards. Dynamic network of proximity interactions for a total duration of 70 days between (A) 43 lizards at site 1, and (B) 44 lizards at site 2. Each temporal slice summarizes interactions within a day (24 hours). Edges indicate that the pair of individuals were within 14m distance of each other, and the edge weights are proportional to the frequency of physical interactions between the node pair. Green nodes are the animals that were diagnosed to be uninfected at that time-point, red are the animals that were diagnosis to be infected and grey nodes are the individuals with unknown infection status at the time-point. We hypothesized that the spatial proximity networks could explain the observed spread of *Salmonella* in the population. The results are summarized as a table. Bold numbers indicate that the network hypothesis was found to have high predictive power compared to an ensemble of randomized networks. The network hypothesis with the highest log Bayesian (marginal) evidence at each site is marked with an asterisk (*).

## Discussion

Network modeling of infectious disease spread is becoming increasingly popular, because modern technology has dramatically improved the quality of data that can be collected from animal populations. However, the concepts of power analysis and hypothesis testing are still underdeveloped in network modeling, even though such approaches are widely recognized as key elements to establish how informative and appropriate a model is (*Jennions and Møller, 2003; Johnson et al., 2015*). Our ability to define a contact network relies on our understanding of host behavior, and the dominant transmission mode for a given pathogen. Since such information is either derived from expert knowledge (which can be subjective) or laboratory experiments (which are time-and resource-intensive), it is essential to conduct an a priori analysis of contact network models to avoid uninformative or misleading disease predictions.

In this study we therefore present INoDS as a tool that performs network model selection and establishes the predictive power of a contact network model to describe the spread of infectious diseases. Our method also provides epidemiological insights about the host-pathogen system by enabling hypothesis testing on different transmission mechanisms, and estimating pathogen transmission rates (transmission parameter, *β*). Unlike previous approaches, our method is robust to missing network data, imperfect disease surveillance, and can provide network inference for a range of disease spread models. The tool can thus be used to provide inference on contact networks for a variety of pathogen types occurring both in wildlife and human populations. Inferring the role of dynamic contacts on infectious disease spread requires the knowledge of either order or timing of infection of individuals in the network. In practice, constraints on data collection (e.g., due to infrequent health assessments), or infection diagnostics (e.g., due to sub-clinical infection, poor diagnostics) precludes precise knowledge of infection timing. To overcome this challenge, our tool assumes the infection times in a host population to be unobserved, and uses data on infection *diagnosis* instead to provide inference on contact networks.

Our work thus addresses a growing subfield in network epidemiology that leverages statistical tools to infer contact networks using all available host and disease data (*Welch et al., 2011; Stack et al., 2013; VanderWaal et al., 2016; Groendyke et al., 2011*). Our approach can be used to tackle several fundamental challenges in the field of infectious disease modeling (*Eames et al., 2015; Pellis et al., 2014*). First, INoDS can be used to perform model selection on contact network models that quantify different transmission modes; this approach therefore facilitates the identification of infection-transmitting contacts and does not rely on laboratory experimentation (or subjective expert knowledge). Second, INoDS can be used to establish the predictive power of proxy measures of contact in cases where limited interaction data is available. For example, spatial proximity, home-range overlap or asynchronous refuge use are commonly used as a proxy for disease-causing contact in wild animal populations (*Godfrey et al., 2010; Leu et al., 2010; Sah et al., 2016*). INoDS establishes the epidemiological significance of such assumptions by comparing the likelihood of infection spread occurring along the edges of the proxy contact network to the likelihoods generated from an ensemble of random networks. Third, it is well known that not all contacts between hosts have the same potential for infection transmission. The heterogeneity of host contacts in a network model is typically captured through edge weights, but it is often not clear which type of edge weights (frequency, duration or intensity) is relevant in the context of a specific host pathogen system (*Pellis et al., 2014*). Through model selection of contact networks with similar edge connection but different edge weighting criterion, INoDS can help establish a link between edge weight and the risk of transmission across an edge in a contact network.

We demonstrate the application of INoDS using two real-world datasets. In the first dataset, we used INoDS to determine the role of edge weight type and edge weight value on the predictive power of the contact network. To accurately model the spread of the *Crithidia* gut protozoan in bumble bee colonies, we show that the contact networks weighted with respect to frequency, rather than duration, have higher predictive power given observed patterns of transmission. Our results therefore support the original finding of the study (*Otterstatter and Thomson, 2007*), where individual risk of infection was found to be correlated with contact rate with infected nest-mates. Our analysis further extends previous findings in this system by comparing the observed patterns of transmission against null expectations from random networks, and assessing the likelihood of competing network hypotheses. We find weak ties below a certain threshold do not play an important role in infection transfer. Contact networks where such weak weighted edges have been removed, therefore, demonstrate higher predictive power than fully connected networks. In the next empirical example, we explore the transmission mechanisms of a commensal bacterium in wild populations of Australian sleepy lizards. We find that taking repeated contacts between closely located lizards into account allows better, i.e. more consistent, predictions on Salmonella transmission.

The current version of INoDS assumes the infection process has no latent period, and that the infectiousness of infected hosts and susceptibility of naive hosts is equal for all individuals in the population. These assumptions can be relaxed to incorporate more complex disease progression. For instance, heterogeneity in infectiousness of infected hosts and the susceptibility of naive hosts can be incorporated as random effects in the model by assuming the two follow a Gaussian distribution. Disease latency can also be incorporated using a data-augmentation technique, similar to what we use for inferring infection times.

Our results show that the data-collection efforts should aim to sample as many individuals in the population as possible, since missing nodes have the greatest impact (rather than missing edges) on the predictive power of network models. Since data-collection for network analysis can be labor-intensive and time-consuming, our approach can be used to make essential decisions on how limited data collection resources should be deployed. Our approach can also be used to improve targeted disease management and control by identifying high-risk behaviors and super-spreaders of a novel pathogen without relying on expensive transmission experiments that take years to resolve.

## Methods

Here we describe INoDS, a computational tool that (*i*) estimates per contact transmission rate (*β*) of infectious disease for empirical contact networks, (*ii*) establishes predictive power of a contact network by comparisons with an ensemble of randomized networks, and (*iii*) enables discrimination of competing contact network hypotheses, including those based on pathogen transmission mode, edge weight criteria and data collection techniques. Two types of data are required as input for INoDS - infection time-series data, which include infection diagnoses (coded as 0 = not infected and 1 = infected), and time-step of diagnosis for all available individuals in the population; and an edge-list of a dynamic (or static) contact network. An edge-list format is simply the list of node pairs (each node pair represents an edge of the network), along with the weight assigned to the interaction, and time-step of interaction, with one node pair per line. The tool can be used for unweighted contact networks - an edge weight of one is assigned to all edges in this case. Time-steps of interactions are not required when analysis is performed on static contact networks. The software is implemented in Python, is platform independent, and is freely available at https://bansallab.github.io/INoDS-model/.

## INoDS formulation

We assume that at each instance the potential of acquiring infection for a susceptible individual *i* depends on the per contact transmission rate *β*, the total strength of interactions with its infected neighbors at the previous time-step, and an error parameter *ε* that captures the force of infection that is not explained by the individual’s social connections. The infection receiving potential, *λ_i_*(*t_i_*), of individual *i* at time *t* is thus calculated as:

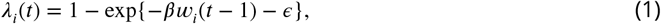

where both transmission rate *β* and error parameter *ε* are > 0; *w_i_*(*t*–1) denotes the total strength of association between the focal individual *i* and its infected associates at the previous time-step (*t* – 1). For binary (unweighted) contact network models *w_i_* = *k_i_*, where *k_i_* is the total infected connections of the focal individual.

The log-likelihood for all observed timings of infection in a population given the contact network hypothesis (*H_A_*) can therefore be estimated as:

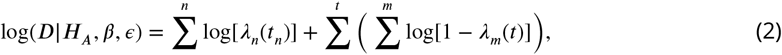

where *t_n_* is the time of infection of individual *n*. The first part of equation 2 estimates the log likelihood of all observed infection acquisition events. The second part of the equation represents the log-likelihood of susceptible individuals *m* remaining uninfected at time *t*.

### Parameter estimation and data augmentation of infection timings

We adopted a Bayesian MCMC framework to estimate the unknown model parameters. Calculation of the likelihood in equation 2 requires knowledge of exact timing of infection, *t*_1_, …*t*_n_, for *n* infected individuals in the population. However in many cases, the only data that is available are the timings of when individuals in a populations were *diagnosed* to be infected, *d*_1_,…*d_n_*. We therefore employ a Bayesian data augmentation approach to estimate the actual infection timings in the disease dataset (*Tanner and Wong, 1987*). Since in this case the actual infection time *t_i_* for an individual *i* is unobserved, we only know that the timing of infection for the individual lies between the interval (*L_i_, d_i_*], where *L_i_* is the last negative diagnosis of individual *i* before infection acquisition. Within this interval, the individual could have potentially acquired infection at any time-step where it was in contact with other individuals in the network. Assuming incubation period to be one time-step, we can therefore represent the potential set of infection timings as *t_i_* ∈ {g_i_ (*t_i_* −1) > 0, *L_i_* <*t_i_* ≤ *d_i_*}, where *g_i_*(*t_i_* − 1) is the degree (number of contacts) of individual *i* at time *t_i_* − 1. For infections that follow a SIS or SIR disease model, it is also essential to impute the recovery time of infected individuals for accurate estimation of infected degree. To do so, we adopt a similar data augmentation approach as described to sample from the set of possible recovery time-points.

The joint posterior distribution of augmented data and the set of parameters is proportional to:

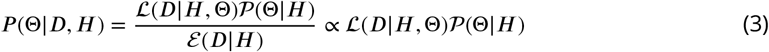

where *D* is the infection time-series data, *H* is the contact network hypothesis, and 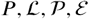 are the shorthands for the posterior, the likelihood, the prior and the evidence, respectively. The data augmentation proceeds in two steps. In the first step, the missing infection times are imputed conditional on the possible set of infection times. In the next step the posterior distributions of the unknown parameters are sampled based on the imputed data. We performed data imputation using inverse transform sampling method, which is a technique of drawing random samples from any probability distribution given its cumulative distribution function (*Robert and Casella, 2004)*. We used a used a uniform prior on [0,1000] for the per contact transmission rate and the error parameter.

MCMC sampling of the unknown parameters is performed using the *PTsampler* function of *emcee* package implemented in Python (*Foreman-Mackey et al., 2013*). *PTsampler* is an implementation of the affine-invariant ensemble MCMC algorithm which provides efficient sampling of highly correlated parameters - a common problem when using simple Metropolis-Hastings type samplers (*Foreman-Mackey et al., 2013*). INoDS uses twice the number of walkers as the total model parameters, and the temperature is set to *T* = 15 to maximize the sampling of the parameter space. The values of The number of sampling steps and burn-in is specified by the user. Convergence is assessed using an autocorrelation plot of few randomly selected walkers. From the joint posterior estimates of *β* and *ε*, we report the parameter combination with the highest maximum likelihood value.

### Statistical significance of infection transmission parameter

The statistical significance of parameter *β* is determined by comparing the force of infection explained by edge connections (=*βw_i_*(*t* – 1)) at each infection event to the error parameter *ε*. The *p*-value is calculated as the proportion of transmission events where the force of infection is greater than the error estimate. The per contact transmission rate *β* is considered to be statistically significant when its calculated *p*-value is less than 0.05.

### Interpretation of the error parameter

In principle, inclusion of the error parameter in eq. 1 is similar to the asocial learning rate used in the network based diffusion analysis approach in the behavior learning literature (*Franz and Nunn, 2009; Aplin et al., 2013*). However, in contrast to the asocial learning rate which quantifies the rate of spontaneous learning, *ε* in INoDS formulation serves to improve the estimation of the per contact transmission rate, *β*, when either the contact network or infection spread is not completely sampled (Appendix Figure 1 and 2). The magnitude of *ε* can also be used to (approximately) assess the magnitude of missing data (Appendix Figure 3). The percentage transmission events where *ε* is greater than the force of infection explained by edge connections (*=βw_i_*(*t* - 1)) increases proportionately with increasing amount of missing network data. The relative differences between social force of infection and unexplained transmission events, however is less sensitive to missing data on infection cases.

### Predictive power of a contact network hypothesis

We assess the predictive power of a contact network hypothesis by performing comparisons with an ensemble of randomized networks with same number of nodes and edge connectivity. Specifically, the likelihood of the infection data given the network and estimated model parameters (i.e., 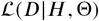) is compared to a distribution of likelihoods of infection data (given the estimated model parameters) obtained from the null networks (i.e., 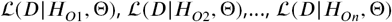); *n* = 500). Null networks are generated by randomizing edge connections of the contact network hypothesis, which preserves the edge density in the permuted networks. Next, a *p*-value is calculated as the proportion of randomizations which generate a likelihood greater than the likelihood of the empirical network hypothesis. The empirical contact network is considered to have a higher predictive power than the null expectation when its calculated *p*-value is less than 0.05.

### Model selection of competing network hypotheses

To facilitate model selection in cases where there are more than one network hypothesis, we compute the marginal likelihood of the infection data given each contact network model. The marginal likelihood, also called the Bayesian evidence, measures the overall model fit, i.e,. to what extent the infection time-series data can be simulated by a network hypothesis (*H_A_*). Bayesian evidence is based on the average model fit, and calculated by integrating the model fit over the entire parameter space:

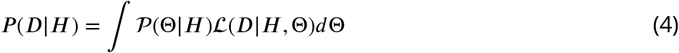

Since it is difficult to integrate Eq.4 numerically, we estimate the marginal likelihood of network models using thermodynamic integration, or path sampling (*Lartillot and Philippe, 2006*) method implemented in *emcee* package in Python. Model selection can be then performed by computing pair-wise Bayes factor, i.e. the ratio of the marginal likelihoods of two network hypotheses. The log Bayes factor to assess the performance of network hypothesis *H_A_* over network hypothesis *H_B_*, is expressed as:

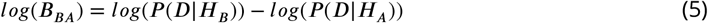

The contact network with a higher marginal likelihood is considered to be more plausible, and a log Bayes’ factor of more than 3 is considered to be a strong support in favor of the alternative network model (*H_B_*) (*Kass and Raftery, 1995*).

### Evaluating INoDS performance

We evaluated the accuracy of the in estimating the unknown transmission parameter *β*, and its robustness to missing data was evaluated. To do so we first constructed a dynamic synthetic network using the following procedure. At time-step *t* = 0, a static network of 100 nodes, mean degree 4, and Poisson degree distribution was generated using the configuration model (*Molloy and Reed, 1995*). At each subsequent time-step, 10% of edge-connections present in the previous time-step were permuted, for a total of 100 time-steps. Next, through the synthetic dynamic network, we performed 10 independent SI disease simulations with per contact rate of infection transmission 0.01 to 0.1. Model accuracy was determined by comparing the estimated transmission parameter, *β*, with the true transmission rate *β^*^* that was used to perform disease simulations. Since the synthetic network dataset did not contain any missing data, model accuracy was also tested by evaluating the deviation of the estimated error parameter *ε*, from the expected value of zero.

We also tested robustness of the tool in establishing the epidemiological relevance of a hypothesized contact network against three potential sources of error in data-collection: missing nodes, missing edges, and missing cases. The three scenarios of missing data were created by randomly removing 10-95% of nodes, edges or infection cases from the simulated dataset described above. True positive rate was calculated as the proportion of times the hypothesized contact network model with missing data was correctly distinguished as statistically significant from an ensemble of null networks generated by randomizing its edge connections. We calculated the true negative rate as the proportion of times a network with the same degree distribution as the contact network hypothesis, but randomized edge connections, was correctly classified as statistically insignificant.

Next, we compared INoDS with two previous approaches (*k*-test and network position test) that have been used to establish an association between infection spread and contact network in a host population. The *k*-test procedure involves estimating the mean infected degree (i.e., number of direct infected contacts) of each infected individual in the network, called the *k*-statistic. The *p*-value in the *k*-test is calculated by comparing the observed *k*-statistic to a distribution of null *k*-statistics which is generated by randomizing the node-labels of infection cases in the network (*VanderWaal et al., 2016*). Network position test compares the degree of infected individuals to that of uninfected individuals (*Godfrey et al., 2009,2010; Maclntosh et al., 2012*). The observed network is considered to be epidemiologically relevant when the difference in average degree between infected and uninfected individuals exceeds (at 5% significance level) the degree difference in an ensemble of randomized networks.

### Applications to empirical data-sets

We demonstrate the applications of our approach using two datasets from the empirical literature. The first dataset comprises of dynamic networks of bee colonies (N = 5-7 individuals), where edges represent direct physical contacts that were recorded using a color-based video tracking software. A bumble bee colony consists of a single queen bee and infertile workers. Here, we focus on the infection experiments in two colonies where infection was artificially introduced through a randomly selected forager (colony UN1 and UN2). Infection progression through the colonies was tracked by daily screening of individual feces, and the infection timing was determined using the knowledge of the rate of replication of *C. bombi* within its host intestine.

The second dataset monitors the spread of the commensal bacterium *Salmonella enterica* in two separate wild populations of the Australian sleepy lizard *Tiliqua rugosa*. The two sites consisted of 43 and 44 individuals respectively, and these represented the vast majority of all resident individuals at the two sites (i.e., no other individuals were encountered during the study period). Individuals were fitted with GPS loggers and their locations were recorded every 10 minutes for 70 days. Salmonella infections were monitored using cloacal swabs on each animal once every 14 days. Consequently, the disease data in this system do not identify the onset of each individual’s infection. We used a SIS (susceptible-infected-susceptible) disease model to reflect the fact that sleepy lizards can be reinfected with salmonella infections. Proximity networks were constructed by assuming a contact between individuals whenever the location of two lizards was recorded to be within 14m distance of each other (*Leu et al., 2010*). The dynamic networks at both sites consisted of 70 static snapshots, with each snapshot summarizing a day of interactions between the lizards. We constructed two contact network hypotheses to explain the spread of salmonella. The first contact network hypothesis placed binary edges between lizards if they were ever within 14m distance from each other during a day. The second contact network assigned edge weights proportional to the number of times two lizards were recorded within 14m distance of each other during a day. Specifically, edge weights between two lizards were equal to their frequency of contacts during a day normalized by the maximum edge weight observed in the dynamic network.

## Acknowledgments

This work was supported by the National Science Foundation Ecology and Evolution of Infectious Diseases Grant 1216054. STL was supported by an Australian Research Council DECRA fellowship (DE170101132), and an ARC grant to CM Bull (DP130100145), which funded the lizard project. We thank Peter Majoros, Tom Haley, Alienor Quiblier and Ben Westwood for their assistance with lizard fieldwork, and David Gordon for his lab work.

## Appendix 1

**Appendix 1 Figure 1.**
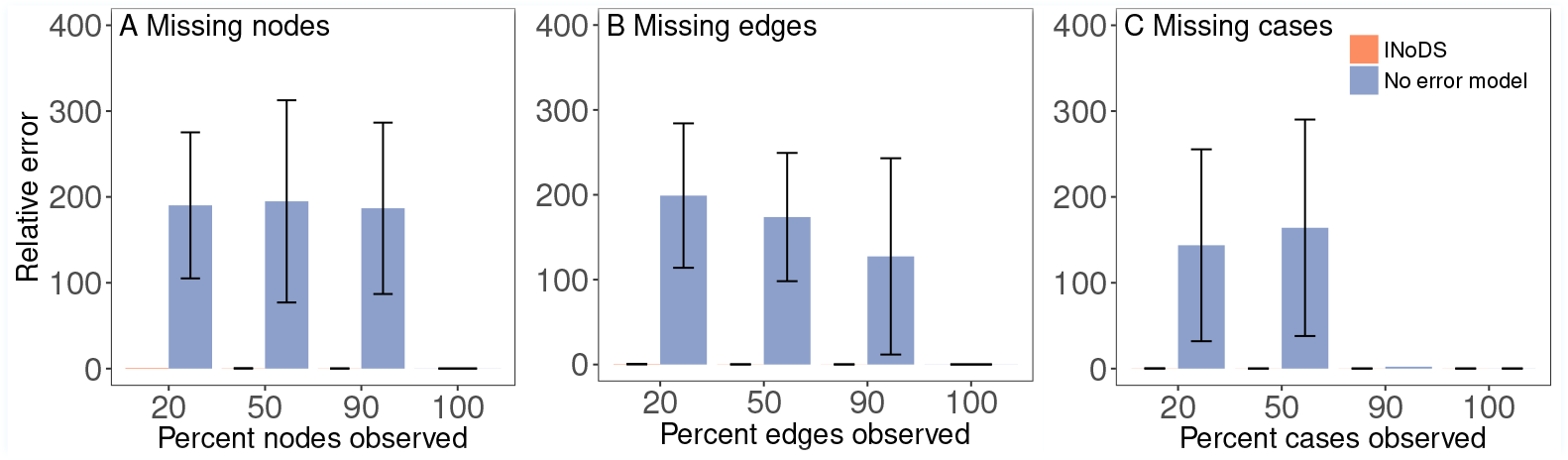
Relative error in the estimations of parameter *β* under missing data conditions with and without the inclusion of the error parameter (*ε*) in the INoDS formulation. The simulated infectious disease spread (SI model, per contact transmission rate pathogen *=* 0.03) was performed on static network with 100 nodes, Poisson degree distribution, and a average degree of 3. Relative error was calculated as 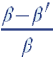, where *β* is the per contact transmission rate of the simulated pathogen (=0.03) and *β’* is the value of social transmission parameter estimated.

**Appendix 1 Figure 2.**
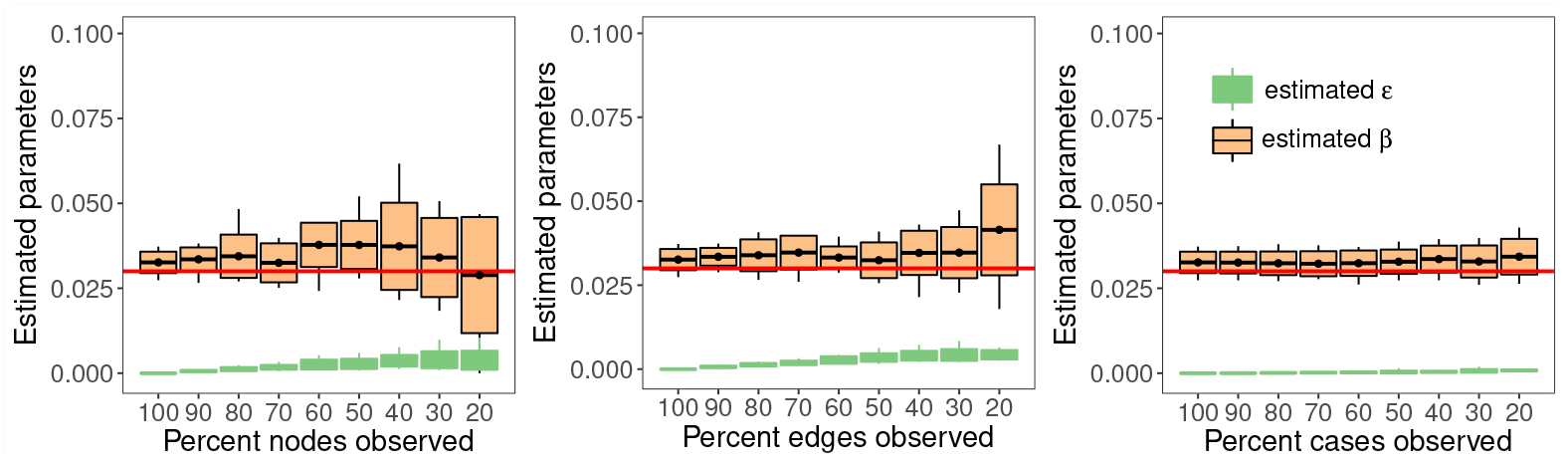
Estimation of per contact transmission rate (*β*) and the error parameter (*ε*) by INoDS under three forms of missing data conditions - (A) missing nodes, (B) missing edges and (C) missing infection cases. Simulations of susceptible-infected (SI) model of infectious disease spread were performed on static network with 100 nodes, Poisson degree distribution, and an average degree of 3. Each boxplot summarizes the results of 10 independent disease simulations; the horizontal line in the middle is the mean of estimated parameter values, the top and the bottom horizontal line is the standard deviation, and the tip of the vertical line represents the maximum/minimum value. The solid red line represents the true value of *β* used in the disease simulations. Since the simulations were performed on a known synthetic network, the expected value of error parameter is zero.

**Appendix 1 Figure 3.**
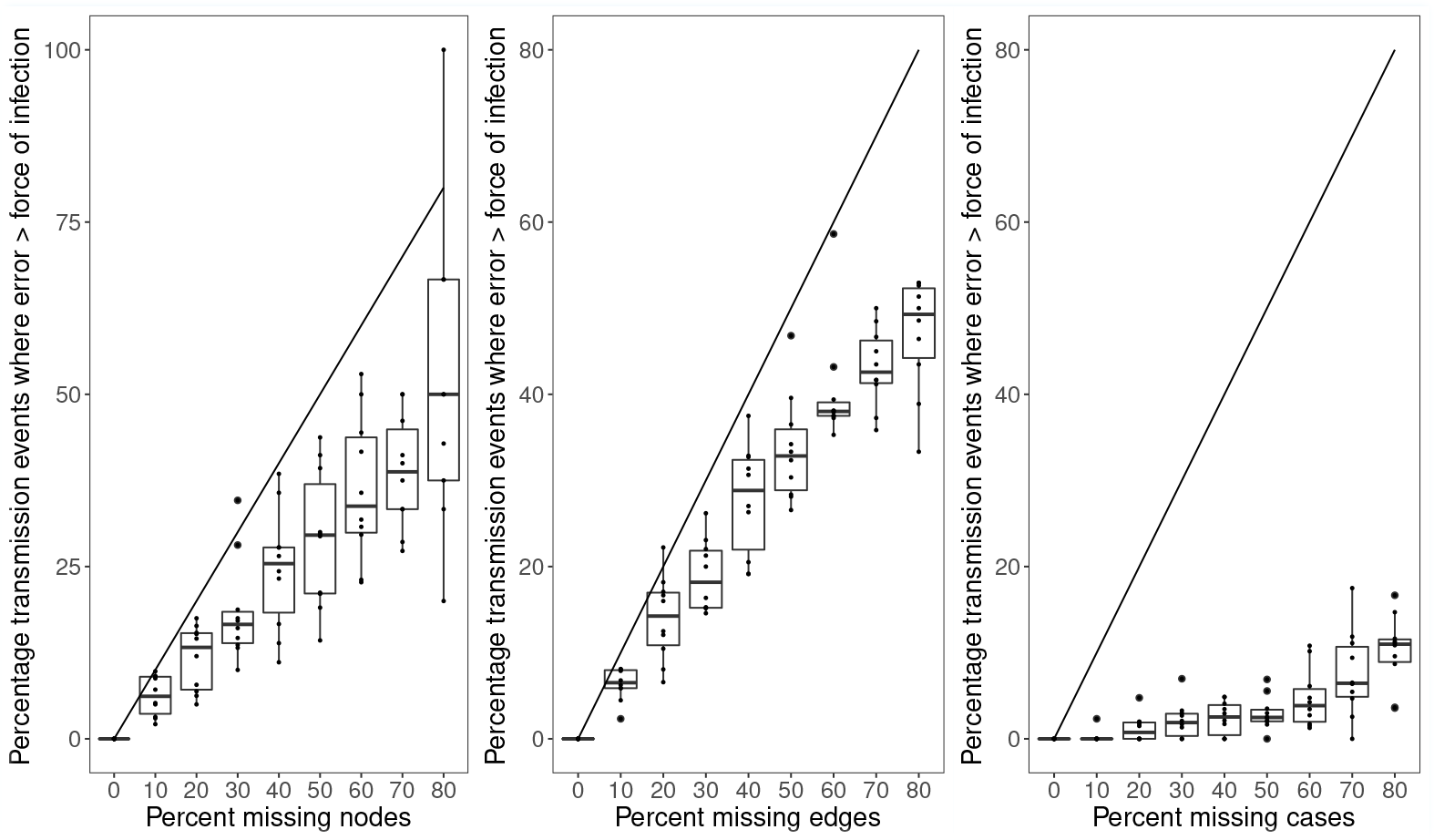
Relationship between error *ε* and force of infection (*=βw_i_*(*t* - 1)) with increasing percentage of missing data. Each boxplot summarizes the results of 10 independent disease simulations (indicated by points); the horizontal line in the middle is the mean percent transmission events where the asocial force is greater than the infection force contributed by the social connections. The top and the bottom horizontal line is the standard deviation, and the tip of the vertical line represents the maximum/minimum value.

**Appendix 1 Figure 4.**
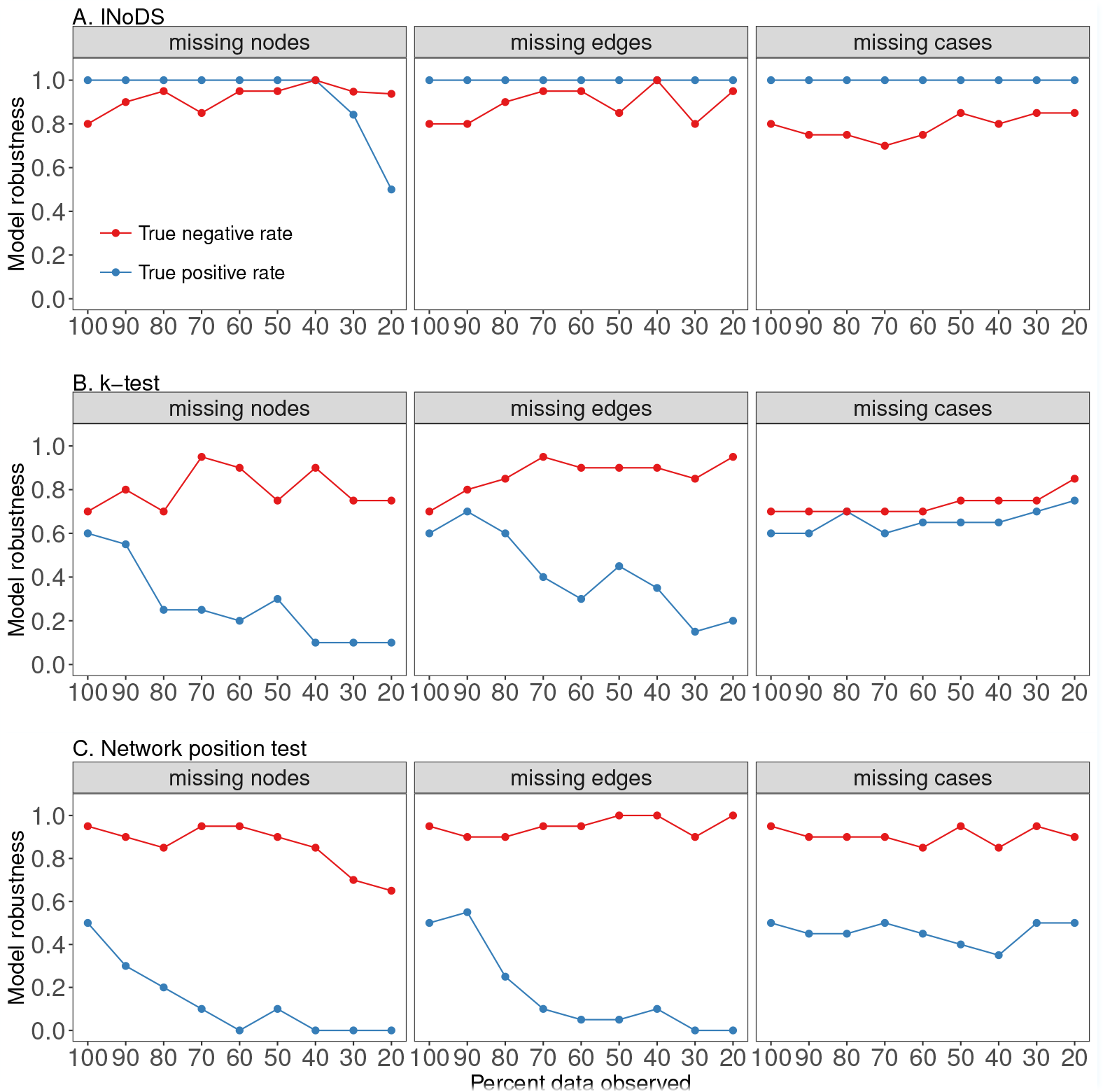
Plot of sensitivity and specificity of (A) INoDS, (B) *k*-test and (C) network position test to three common forms of missing data - missing nodes, missing edges and missing infection cases. The observed network in this case is a static network with 100 nodes, Poisson degree distribution and a mean network degree of 3. Simulations of pathogen spread with per contact transmission rate of 0.03 were performed through the observed static network. Null expectation in INoDS and network position test was generated by permuting the edge connections of the observed networks, creating an ensemble of null networks. In *k*-test, the location of infection cases within the observed network are permuted, creating a permuted distribution of *k*-statistic (VanderWaal et al., 2016).

**Appendix 1 Figure 5.**
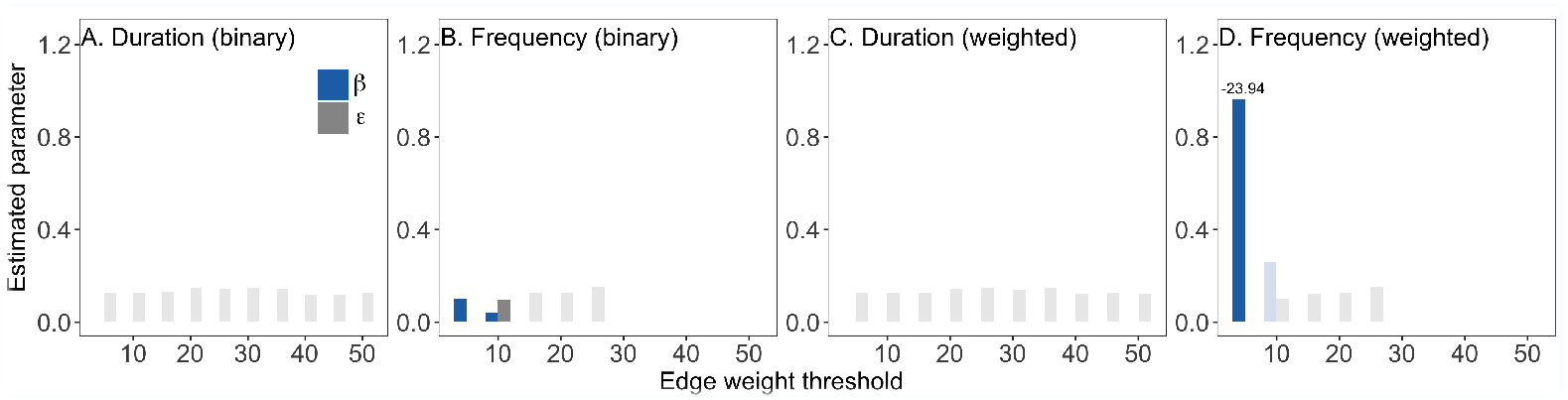
Identifying the contact network model of *Crithidia* spread in bumble bee colony (colony UN2). Edges in the contact network models represent physical interaction between the bees. Since the networks were fully connected, a series of 1ltered contact networks were constructed by removing weak weighted edges in the network. The x-axis represents the edge weight threshold that was used to remove weak edges in the network. Two types of edge weights were tested - duration and frequency of contacts. In addition, across all ranges of edge weight threshold, the weighted networks were converted to binary networks. The results shown are estimated values of the per contact rate of infection transmission *β*, and estimated values of error *ε*, for the (A-B) two types of binary network, (C) contact duration weighted network, (D) contact frequency weighted network. The faded bars correspond to networks where *β* parameter is statistically insignificant. Numbers above bars indicate the log Bayesian (marginal) evidence of the networks that were detected to have statistically signi1cant higher predictive power as compared to an ensemble of null networks (P < 0.05, corrected for multiple comparisons). We note that frequency networks with more than 25% weak edge removed failed to converge in (C) and (D), and therefore the transmission parameter associated with these contact networks were not estimated.

---

**Figure 3–Figure supplement 1.**
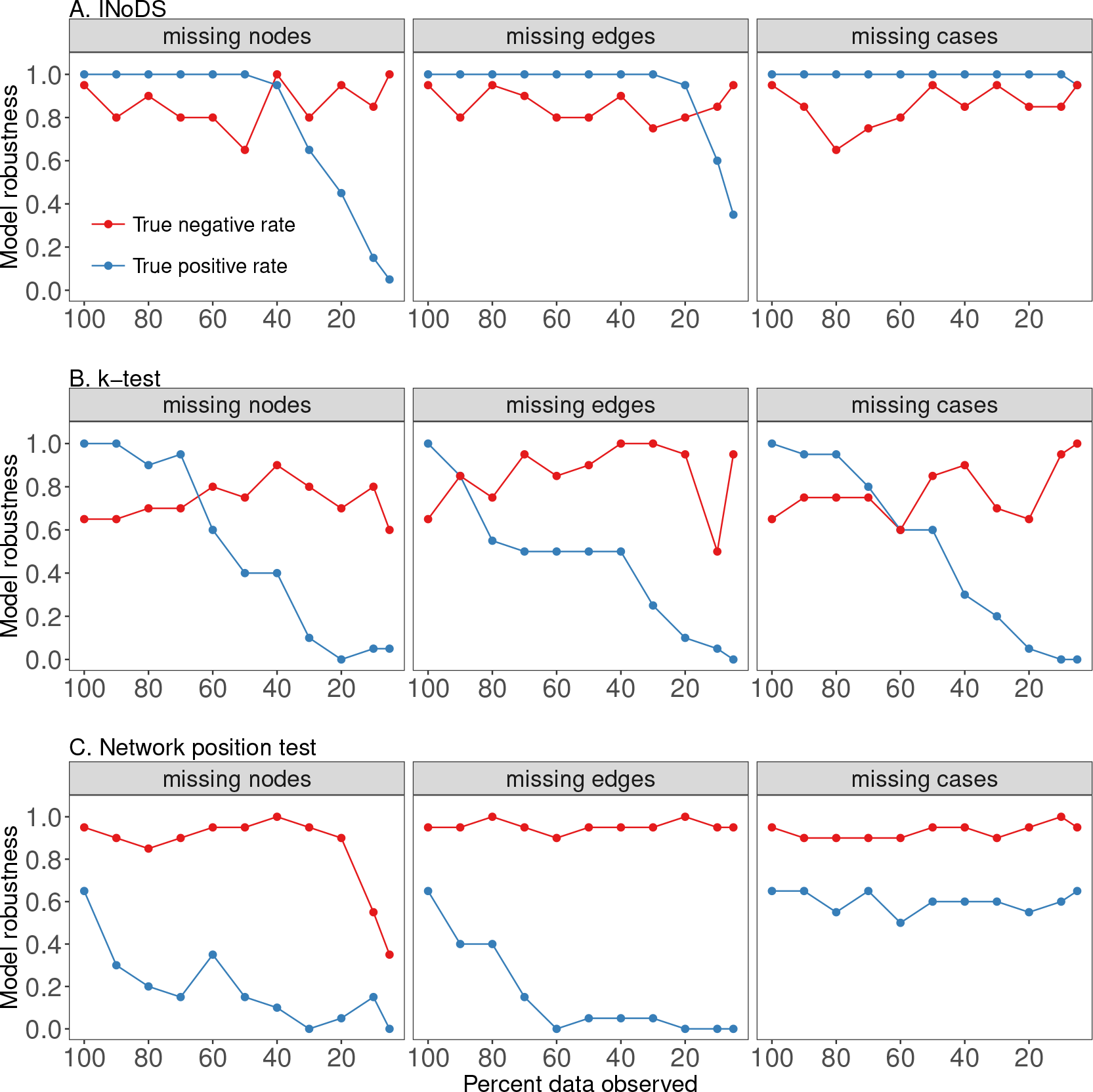
Robustness plot of (A) INoDS, (B) *k*-test and (C) network position test to three common forms of missing data - missing nodes, missing edges and missing infection cases. The null expectation in INoDS and the network position test was generated by permuting network edges, creating an ensemble of null networks. In the *k*-test, the location of infection cases within the observed network are permuted, creating a permuted distribution of the *k*-statistic (*VanderWaal et al., 2016*).

